# Anatomical and cellular heterogeneity in the mouse oviduct-- its potential roles in reproduction and preimplantation development

**DOI:** 10.1101/2020.08.23.263772

**Authors:** Keerthana Harwalkar, Matthew J Ford, Katie Teng, Nobuko Yamanaka, Brenna Yang, Ingo Burtscher, Heiko Lickert, Yojiro Yamanaka

## Abstract

The oviduct/fallopian tube is a tube-like structure that extends from the uterus to the ovary. It is an essential reproductive tissue that provides an environment for internal fertilization and preimplantation development. However, our knowledge of its regional and cellular heterogeneity is still limited. Here, we examined the anatomical complexity of mouse oviducts using modern imaging techniques and fluorescence reporter lines. We found that there are basic coiling patterns and turning points in the coiled mouse oviduct can serve as reliable landmarks for luminal morphological regionalities. We identified previously unrecognized anatomical structures in the isthmus and uterotubal junction (UTJ) that likely play important roles in reproduction. Interestingly, during ovulation, the isthmus was transiently plugged by a thick mucus, keeping the oocytes within the ampulla. Preimplantation embryos travelled along the oviduct and formed a queue within small compartments of the UTJ before uterine entry. Taken together, the oviduct luminal epithelium had highly diverse luminal structures with distinct cell populations reflecting its complex functions in reproduction.

## INTRODUCTION

Mouse oviducts, called fallopian tubes in humans, are a part of the female reproductive tract, forming a tube-like structure that extends from the uterus to the ovary. This tissue is essential for mammalian reproduction, including internal fertilization and preimplantation development. The two gametes, sperm and oocytes, enter from opposite ends of the oviduct to meet at a swollen sac-like region of the oviduct, the ampulla (AMP), for fertilization. After fertilization, preimplantation embryos travel down the oviduct lumen toward the uterus (Kölle et al., 2020; Moore et al., 2019). They remain in the oviduct for relatively consistent periods of time without direct physical contact between the embryos and oviduct luminal surface due to the zona pellucida, a shell-like structure preventing their physical interaction (Wassarman et al., 1999). The oviduct luminal environment is transiently modulated by oviduct luminal epithelial cells to accommodate sperm migration, fertilization, and preimplantation development during the estrus stage (Barton et al., 2020; Li & Winuthayanon, 2017; Winuthayanon et al., 2015). Without this regulation, the oviductal luminal environment is too harsh for reproduction because it is primarily adjusted to prevent bacterial infection. On the other hand, if a preimplantation embryo does not properly travel the tube, for example, due to an obstruction or constriction in the luminal space, this leads to exo-uterine pregnancy in humans (Marion & Meeks, 2012). This indicates that a proper spatiotemporal coordination between the oviduct and gamete/embryo movement is essential for successful pregnancy.

Although this tissue is essential for reproduction, with a portion of human infertility cases being due to deficiencies in this tissue (Briceag et al., 2015), the tissue structure and cellular heterogeneity of the oviduct have not been fully explored. The pioneer work on the mouse oviduct (Adguhr, 1927) described intensive oviduct coiling and complex luminal morphology. Four regionalities are recognized in the mouse oviduct, from the distal end: the infundibulum (INF), ampulla (AMP), isthmus (ISM) and uterotubal junction (UTJ). It has been generally thought that the oviduct luminal epithelium consists of two cell types: multi-ciliated and non-ciliated/secretory cells, with the proportion of multi-ciliated cells forming a descending gradient from the distal to proximal (Dirksen, 1972; Stewart & Behringer, 2012). However, in our parallel study, we identified that distal and proximal luminal epithelial cells are two distinct lineages from as early as E12.5 in the Mullerian duct with unique gene expression (Ford et al., 2020).

In this study, we revisit this essential tissue using modern imaging techniques and mouse fluorescence reporter lines. Due to its luminal epithelial folds and intensive coiling, traditional 2D tissue sectioning has limitations in analysis of 3D complexity. Through our careful 3D analysis, we recognized basic coiling patterns, coinciding with changes in luminal regional morphology. In addition, we identified previously unrecognized anatomical structures in the ISM and UTJ, likely playing important roles for successful pregnancy. The AMP-ISM junction (AIJ) was a unique regionality different from the AMP and ISM in its multi-ciliated cell distribution pattern, luminal morphology, transcriptional factor expression, and acidic mucin secretion. Our results revealed anatomical and cellular heterogeneity in the mouse oviduct luminal epithelium, suggesting functional diversity in each morphologically distinct region.

## RESULTS

### Basic coiling patterns in mouse oviducts maintained by its attachment to the mesosalpinx

The mouse oviduct is a highly coiled tube that extends from the uterus to the ovary. This coiling makes precise regional comparison difficult in standard 2D tissue sectioning. The oviduct mesosalpinx, a thin layer of mesentery, gathers behind the coils to support the turning points. It connects to the ovarian bursa, a thin membrane that encompasses the ovary, as well as the uterine mesometrium, which is further connected to the dorsal mesentery (Adguhr, 1927). We carefully compared several female reproductive tracts (n=10 oviducts) to examine if they have basic coiling patterns (Fig.1A,B; Suppl. Fig.1). Although occasional contractions of the AMP and ISM myosalpinx created variation (Dixon et al., 2009), their basic coiling patterns appeared consistent. The coiling pattern of the left and right oviducts were mirror images of each other. When the right oviduct was placed with the ovary on top and the uterus at bottom, the distal AMP was always positioned to the left side and the proximal ISM was positioned to the right side (Fig.1A; Suppl. Fig.1).

Based on the basic coiling patterns, we numbered each turn from the distal end (Fig.1C). Turns 1 to 4 were highly consistent with Turn 4 often hidden under Turns 2 and 3. Turns 5 to 7 were more varied likely due to ISM contractions and Turn 5 frequently curled inward (Fig. 1D) creating additional Turns 4.5 and 5.5. Turns 9-11 were again relatively consistent with Turn 11 often lying behind Turns 9 and 10 and connected to the uterus. When the oviduct was straightened, each turn was still recognizable and its luminal fold patterns along the oviduct were visible under a dark field illumination of a dissecting microscope (Fig.1E). The turning positions were reliable landmarks for luminal fold patterns along the mouse oviduct.

**Figure 1:**
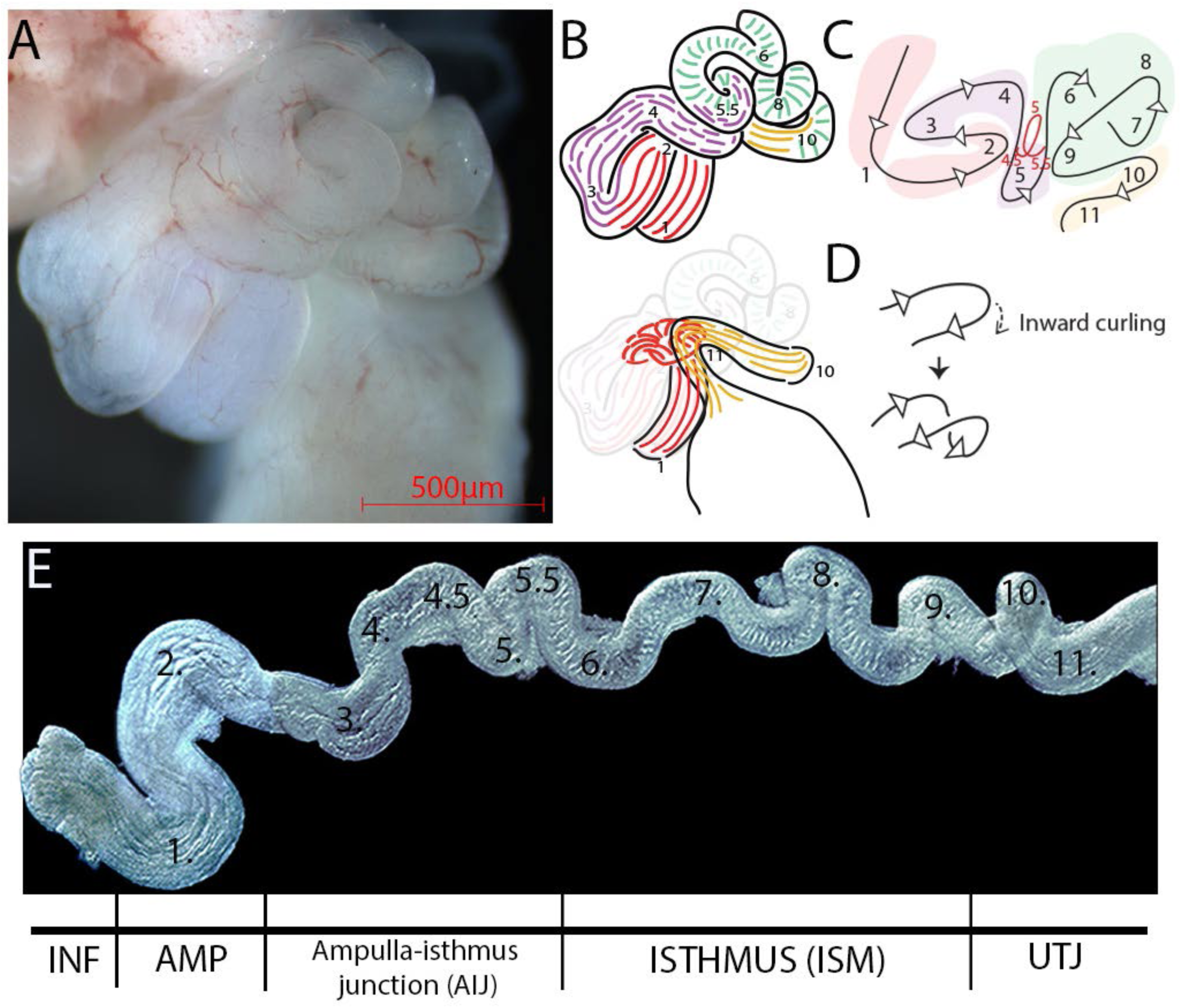
Basic coiling pattern and turning points in the mouse oviduct. (A) The dorsal view of the right oviduct, with AMP on the left side. (B) The diagrams of the oviduct shown in A. Surface view (top). Perspective view (bottom). The INF and UTJ were hidden under the body of the oviduct. (C) A simple illustration of a basic coiling pattern and regionalities. 11 turns were reproducibly identified. Turn5 was often curled inward (red curve), creating Turn 4.5 and 5.5. The coiling pattern from Turn5 to Turn7 showed variation due to ISM contractile movement. The areas of Turn1-3 and Turn8-11 had consistent coiling pattern. The colours represent distinct luminal fold morphologies. (D) Basic coiling pattern in the ISM. The complex coiling pattern is created by a combination of turning and curling. (E) A stretched oviduct with mesosalpinx removed. Tiled images of the stretched oviduct were manually aligned. A plain black background was added. N=10 oviducts.

### 3D imaging reveals transition of epithelial fold patterns along the mouse oviduct

In order to investigate luminal epithelial fold patterns and their transition along the oviduct, we undertook high resolution 3D confocal imaging analysis after tissue clearing. The INF is a funnel-like structure located at the distal end of the oviduct, adjacent to the ovary (Agduhr, 1927). Similar to the collar of a turtleneck sweater, the interior luminal mucosal epithelium of the INF folded over the exterior (Fig.2A,D,G), exposing the mucosal epithelium to the inner space of the ovarian bursa (Suppl. Fig.2). At the opening of the oviduct, called the ostium, almost every two exterior longitudinal folds merged into one luminal fold and continued into the AMP (Fig.2E). The end of the oviduct was often beveled to fit the ostium along the ovarian surface, with a slit at one side adjacent to the ovary (Fig.2G, Video 1). In contrast to the circular smooth muscle layer, the exterior folds framing the ostium were not radially symmetric (Fig.2B,D, Suppl. Fig.3A). The exterior folds on the lateral sides of the ostium were taller and complex, making the relatively flat wide surface at the beveled end (Fig.2G, Video 1). The marginal edges of these tall lateral exterior folds were not attached to the tube but formed an overhang (Fig.2D,F, Video 1). Behind the ostium, the exterior folds were shorter and laid flat against the smooth muscle layer (Fig.2B,F, Video1). The mucosal epithelium of these exterior folds was continuous with the epithelium-lined ligament that connected to the ovarian bursa and ovary (Fig.2F, Suppl. Fig.2, 3A). The INF luminal folds consisted of 12-18 longitudinal folds (Fig.2B, Suppl. Fig.3A; n=8 oviducts). Every other luminal fold was tall (80-184um), almost reaching the center of the lumen, with a short fold roughly half the height (16-80um) between two tall ones (Fig. 2B, Suppl. Fig. 3A). Each fold has a thin layer of stromal cells tacked inside without smooth muscle cells. The smooth muscle layer is only at the circumference, framing the tube-like shape at the INF (Agduhr, 1927).

**Figure 2:**
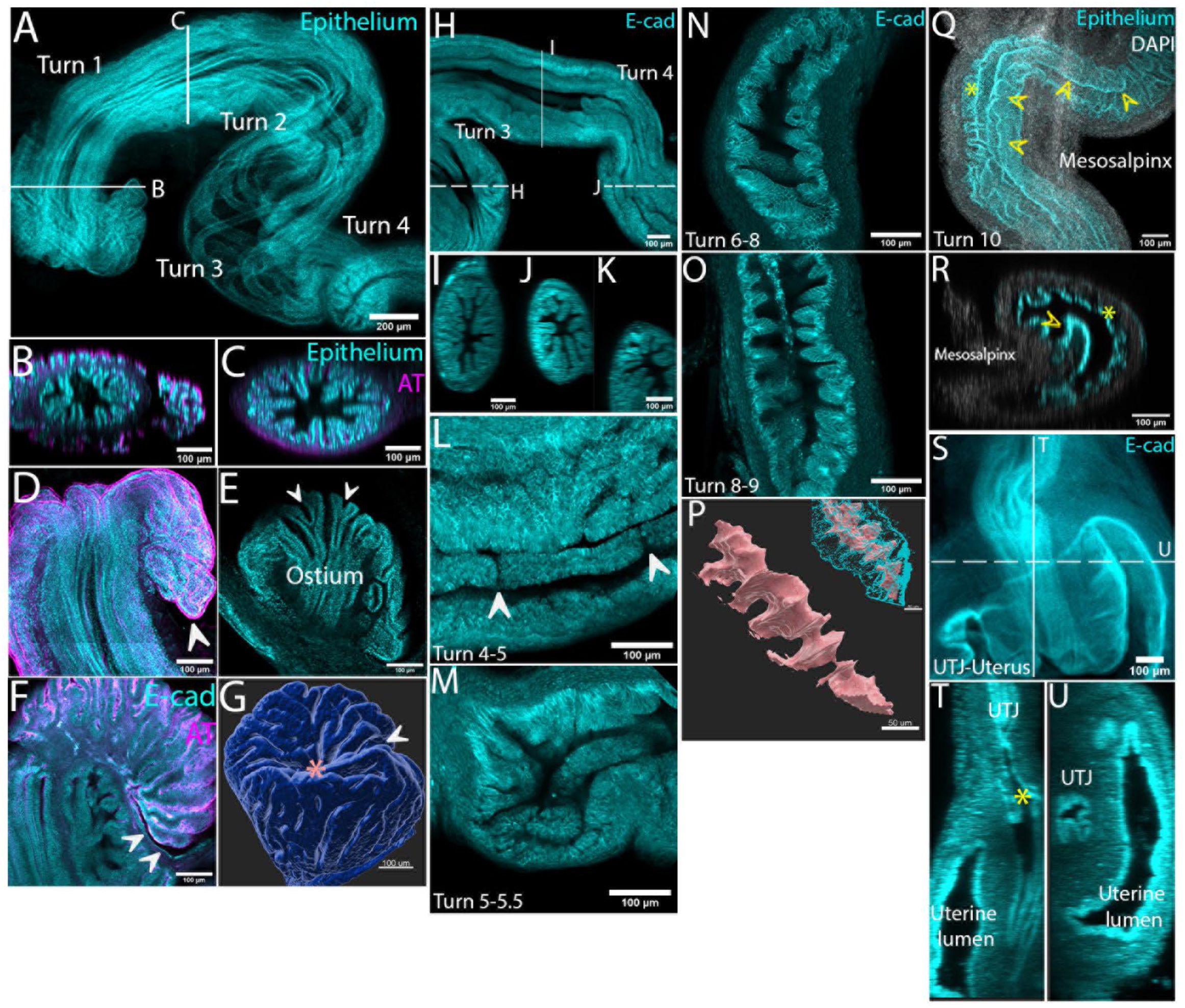
Distinct 3D luminal morphologies along the mouse oviduct. (A) The distal end of the oviduct from Turn 1 to 4, including the INF, AMP and AIJ. Continuous, longitudinal folds extended from the INF into Turn 3-4. Sum slices projection of 90 optical sections. White lines indicate location of orthogonal views in (B) and (C). (B) XZ orthogonal view of the INF. (C) YZ orthogonal view of the AMP. Tall and short luminal folds alternate along the circumference. (D) Exterior folds of the INF were continuous with the luminal folds. The marginal edge of the exterior folds formed an overhang (white arrowhead). Maximum projection of 23 optical sections. (E) Two exterior folds merged into a single luminal fold at the ostium (white arrowheads). (F) The backside view of the ostium. The mucosal epithelium of overhanging exterior folds was connected to an epithelium lining the ovarian bursa (white arrowheads). (G) Surface rendered image of the ostium. The exterior folds created a relatively wide flat surface lading to the ostium (pink asterisk). A slit opening towards the ovary (white arrow). (H) Continuous longitudinal folds from Turn 3 to 4 of the AIJ. White lines indicate the location of orthogonal views in I-K. (I) The area between Turn 2-3. (J) Turn 3-4 (K) Turn 4-5. A decrease of the number of longitudinal folds was noted. (L) The distal AIJ at Turn 4-5. Breakpoints in the longitudinal folds (white arrows). (M) The proximal AIJ at Turn 5-6. Short and bent longitudinal folds and luminal convolutions. (N) The distal ISM at Turn 6-8. Transverse luminal folds were angled. (O) The proximal ISM at Turn 8-9. Aligned, parallel transverse luminal folds. (P) Surface rendering of the bellows-shaped isthmic lumen. A merged image of the epithelial cell surface rendering and luminal surface rendering (inset). Scale bar = 50 µm (Q) The transition between the ISM and UTJ around Turn 10. One longitudinal fold (yellow arrowheads) on the mesosalpinx side extended into the UTJ, with isthmic transverse folds on the anti-mesosalpinx side (yellow asterisk). (R) XZ orthogonal view in (Q). (S) The UTJ-uterus connection. The UTJ lumen, located within the uterine stroma, connected into the uterine lumen. Average projection of 115 optical sections. (T) Longitudinal orthogonal view in (S). Breaking points in the longitudinal folds formed small compartments (yellow asterisk). (U) Transverse view in (S). Intramural UTJ was at the mesosalpinx side of the uterine lumen. Epithelium: a combined signal of PAX8 and FLTP-H2B-Venus (used in 2A-E, Q, R; see M&M). AT: acetylated tubulin. E-cad: E-cadherin. N=4 mice. Scale bars = 100 µm unless mentioned above.

**Figure 3:**
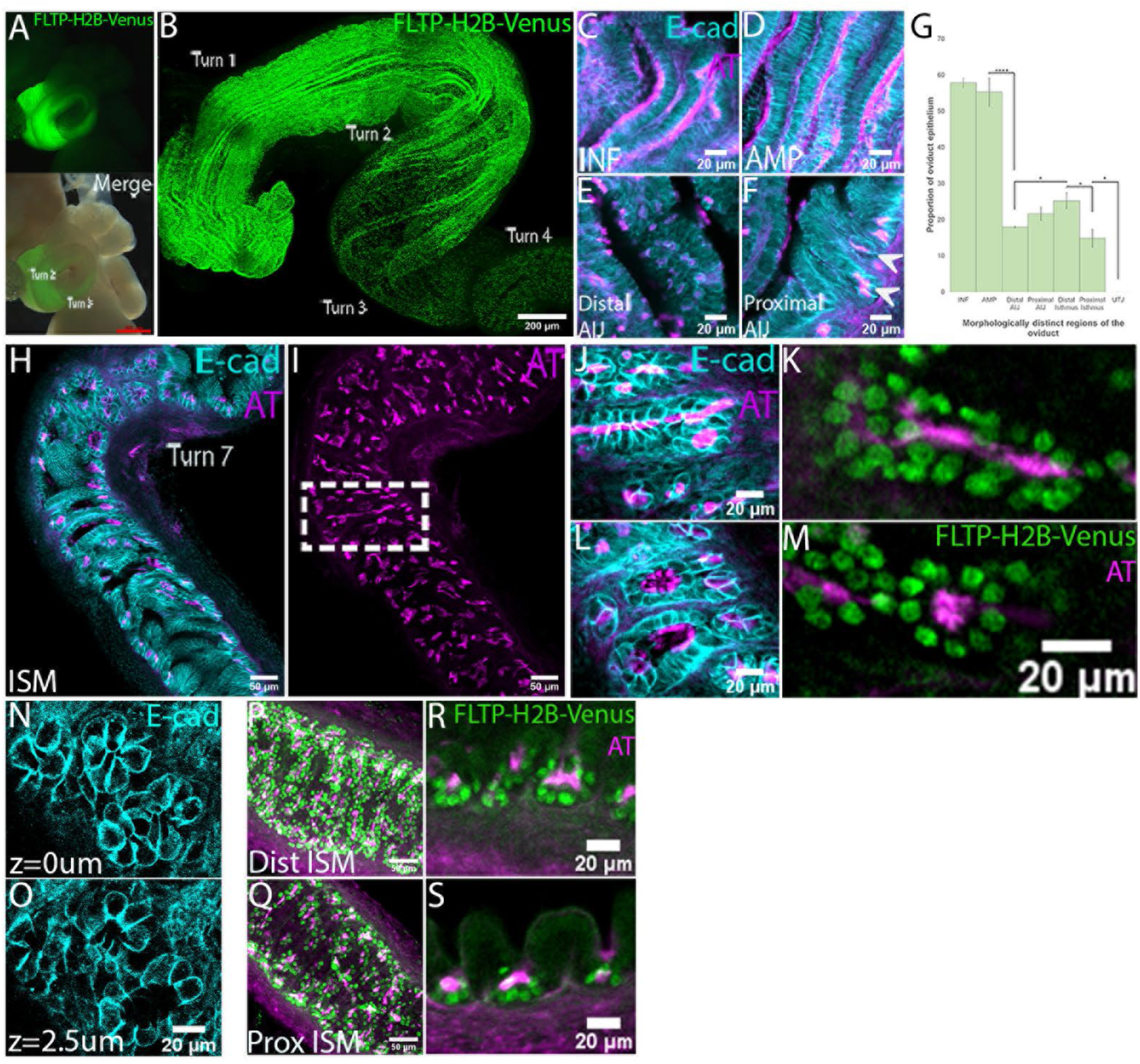
Distribution of multi-ciliated cells in the oviduct and unique multi-ciliated cell clusters in the ISM. (A) An oviduct of a FLTP-H2B-Venus female. Strong GFP signal in the INF and AMP. Scale bar = 500 µm. (B) Distribution of FLTP-H2B-Venus cells in the INF, AMP and AIJ. A sharp reduction of H2B-Venus+ve cells between Turn 2 and 3. The same image series as Fig.2A. Maximum projection of 90 optical sections. Scale bar = 200 µm (C-F) Distribution patterns of multi-ciliated cells in the INF, AMP and AIJ. (C, D) Uniform distribution of multi-ciliated cells in the INF (C) and AMP (D). Cilia are evenly distributed on the apical surface. (E) Multi-ciliated cells are sparsely distributed in the epithelium of the distal AIJ. Cilia localization is not uniform but often limited at the periphery or center of the apical surface of multi-ciliated cells. (F) Multi-ciliated cells in the proximal AIJ. They are located in the trenches (white arrowheads), similar to the ISM, but some are sparsely distributed on the folds. (G) Proportions of FLTP-H2B-Venus+ve multi-ciliated cells along the oviduct. Regions of high (around 60% in INF/AMP), low (around 20% in AIJ/ISM) and absent (0% in UTJ) are recognized. ****<0.0001, *<0.05. Quantification was performed on transverse sections like Suppl.Fig.2. N= 8 oviducts. (H-S) Multi-ciliated cells in the ISM. (H) A single optical section of the distal ISM. All multi-ciliated cells are clustered in the trenches of the epithelial folds. (I) Maximum projection of acetylated tubulin staining from 41 optical sections. A stripe pattern of the multi-cilia distribution, resembling the sperm distribution pattern (La Spina et al., 2016; Muro et al., 2016). (J,K) A groove of multi-ciliated cell clusters. (L,M) Multi-ciliated cell pits. Cilia project to one focal point. (N, O) A rosette arrangement of multi-ciliated cell clusters forming grooves and pits. Bottom of pits (N). Close to luminal surface (O). (P,Q) Difference in density of multi-ciliated cells along the ISM. Max projections of FLTP-H2B-Venus and AT staining of the distal (P) and proximal (Q). Higher density of multi-ciliated cells in the distal ISM than in the proximal ISM. (R,S) Longitudinal orthogonal views of distal (R) and proximal (S) isthmic transverse folds. More multi-ciliated cells contribute to form single pits/grooves in the distal ISM. Scale bars = 20 µm/ 50 µm.

The longitudinal luminal folds of the INF were continuous beyond Turn1, leading into the AMP (Fig.2A). The AMP had 15-20 inner luminal folds (Fig.2C, Suppl. Fig.3B; n=8 oviducts). These longitudinal folds became fewer after Turn 3 (5-12 luminal folds; Fig.2H-K, Suppl. Fig.3C,D; n=8 oviducts). In the AMP-ISM junction (AIJ), between Turns 4-5.5, the longitudinal folds had breaking points (Fig.2L). Towards the ISM, the longitudinal folds were often bent and had frequent breaking points with multiple convolutions (Fig.2M). The ISM started around Turn 6, with transverse luminal folds (Fig.2N) forming a bellows-shaped lumen (Fig.2P). In the distal ISM ranging from Turn 6-8, the folds were not well aligned but often fused and angled at each other (Fig.2N), while in the proximal ISM ranging from Turn 8-9, they were relatively aligned and parallel (Fig.2O).

The transition to the UTJ was around Turn 10 with the formation of continuous longitudinal luminal folds (Fig.2Q). In this transition area, the isthmic transverse folds were on the anti-mesosalpinx side while 1-2 longitudinal folds extended from the UTJ on the mesosalpinx side (Fig.2Q,R). At the junction where the oviduct lumen is inserted to the uterus, called the intramural UTJ (Fig.2S), all longitudinal folds simultaneously had breaking points, creating small compartments (Fig.2T, Video 2). The intramural UTJ lumen was connected to the uterine lumen through the colliculus tubarius (Fig. 2U), as described previously (Agduhr, 1927; Muro et al., 2016).

### Multi-ciliated cells are clustered in epithelial trenches of the isthmus

It is thought that the proportion of multi-ciliated cells forms a descending gradient from the distal end (Dirksen, 1972; Stewart & Behringer, 2012). Since the mouse oviduct has complex luminal folds and a highly coiled structure, it is important to perform 3D image analysis to capture the whole view of their distribution pattern. Using a FLTP-H2B-Venus transgenic mouse line (Gegg et al., 2014) for visualisation of multi-ciliated cells, we found that the distribution of FLTP-H2B-Venus+ve cells did not show a simple gradient but a sharp reduction between Turn 2 and 3 (Fig.3A,B). A high proportion of FLTP-H2B-Venus+ve cells in the INF and AMP was observed (58% and 55%, respectively; Fig.3G, Suppl. Fig.3A,B), with a relatively low proportion in the AIJ and ISM (ranging from 18 to 25%; Fig.3G, Suppl. Fig.3C-F). No FLTP-H2B-Venus+ve cells were found in the UTJ (Suppl. Fig.3G).

In the INF and AMP, multi-ciliated cells were uniformly distributed on the epithelial folds (Fig.3C,D). In the distal AIJ around Turn 3, they were sparsely distributed (Fig.3E). Some multi-ciliated cells in the proximal AIJ were located in the trenches of transverse luminal folds, similar to the ISM (see below), while others were sparsely distributed on the folds (Fig.3F). Although the location of multi-ciliated cells changed progressively, there was no significant difference in the proportion of multi-ciliated cells within the AIJ (18% in distal compared to 19% in proximal, Fig.3G).

Interestingly, in the ISM, all multi-ciliated cells clustered together in the trenches of the transverse folds (Fig.3H). They formed a rosette arrangement (Fig.3N,O) that generated grooves and pits of multi-ciliated cells (Fig.3J-M). All cilia were projected into one focal point within the grooves and pits (Fig.3J-M), creating a stripe pattern of the cilia distribution in the ISM (Fig.3I). This stripe pattern is very similar to the sperm distribution pattern in the mouse oviduct in previous studies (La Spina et al., 2016; Muro et al., 2016). Although the trench localization of multi-ciliated cells in both distal and proximal ISM did not change (Fig.3P-S), there was a significant decrease in the proportion of multi-ciliated cells in the proximal ISM (distal 25% and proximal 15%; Fig.3G). The density of multi-ciliated cells was lower in the proximal ISM (Fig.3P,Q) because the number of multi-ciliated cells forming each pit or groove was smaller (Fig.3Q,S).

### Distinct subtypes of secretory and multi-ciliated cells in distal and proximal regions in mice and marmosets

In a parallel study, we demonstrated that distal and proximal oviduct luminal epithelial populations are two developmentally distinct lineages from as early as the E12.5 Mullerian duct. Based on single cell transcriptome analysis, we identified that WT1 and PAX2 are specific markers for the distal and proximal regions, respectively. Interestingly, PAX8 is a marker for secretory cells in the distal region but is uniformly expressed in all epithelial cells, secretory and multi-ciliated cells, of the proximal oviduct epithelium (Ford et al., 2020; Fig.4A-F, Suppl. Fig.4A,B).

We searched for other transcription factors differentially expressed in the two regions and identified SOX17 and FOXA2. SOX17 and FOXA2 are both expressed in the uterine glandular epithelium but not in the uterine luminal epithelium (Hirate et al., 2016; Spencer et al., 2017). In contrast to their co-expression pattern in the uterine gland, their expression patterns in the oviduct were different. The expression of SOX17 in the oviduct was similar to PAX8 (Fig.4G-L, Suppl. Fig.4C,D). SOX17 was expressed in most of PAX8+ve non-ciliated secretory cells of the INF and AMP (82% and 72% of PAX8+ve secretory cells, respectively). Some double negative non-ciliated cells (PAX8-ve; SOX17-ve) were also present (10% in INF and 11% in AMP). On the other hand and identical to PAX8, SOX17 was uniformly expressed in the proximal regions (Fig.4H-L, Suppl. Fig.4C,D). Interestingly, FOXA2 was uniquely expressed in multi-ciliated cells of the INF and AMP, but absent from multi-ciliated cells of the ISM (Fig.4M-R, Suppl. Fig.4E,F).

**Figure 4:**
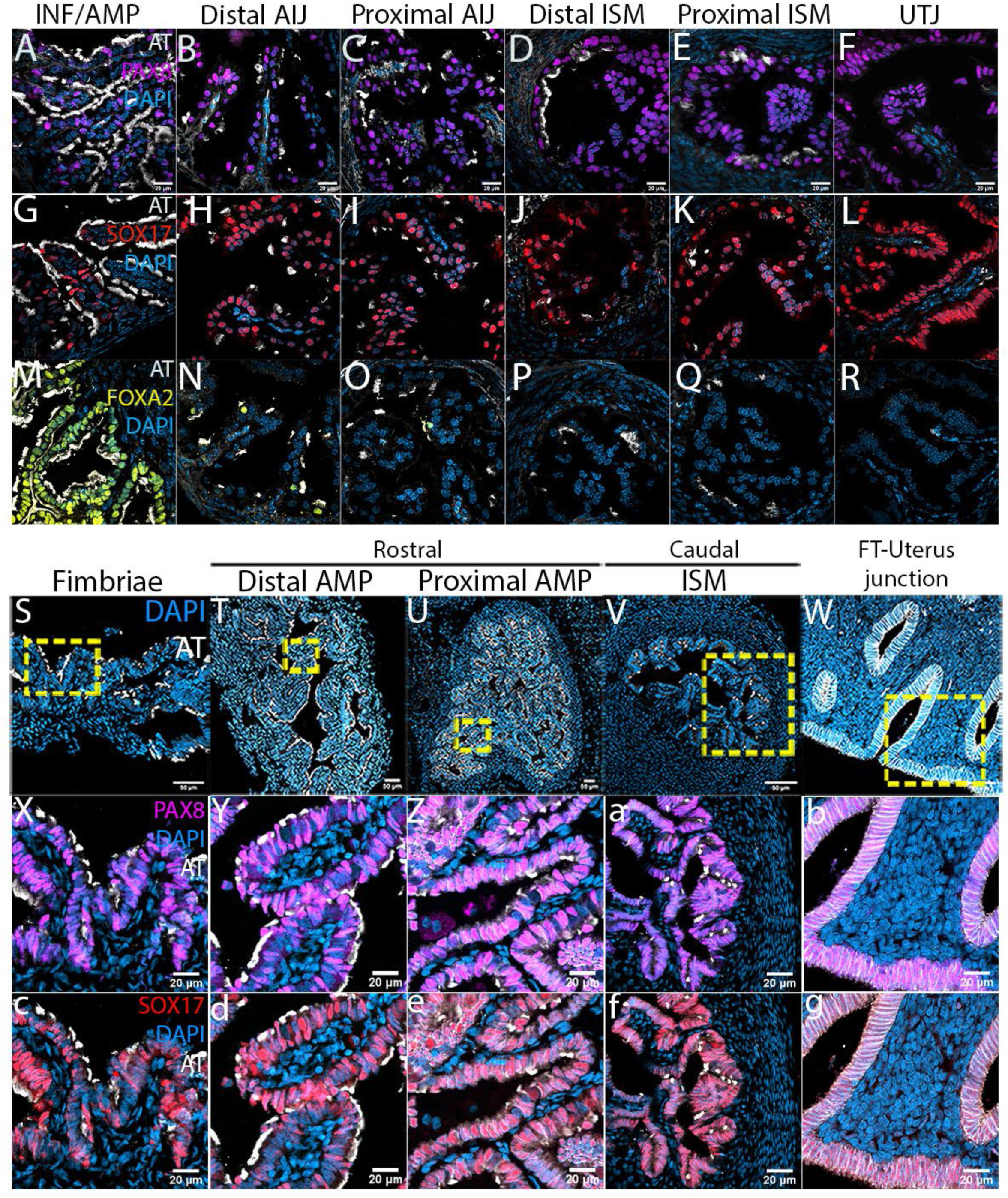
Distinct expression patterns of transcription factors in distal and proximal luminal epithelium in mice and marmosets. Expression patterns of PAX8 (A-F), SOX17 (G-L), and FOXA2 (M-R) in the mouse oviduct. (A,G,M) INF/AMP. (B,H,N) Distal AIJ. (C,I,O) Proximal AIJ. (D,J,P) Distal ISM. (E,K,Q) Proximal ISM. (F,L,R) UTJ. (A,G) PAX8 and SOX17 were expressed in non-ciliated secretory cells in the INF and AMP. (B-F,H-L) PAX8 and SOX17 were expressed in all epithelial cells, both multi-ciliated and non-ciliated secretory cells. (M) FOXA2 was expressed in multi-ciliated cells in the INF and AMP. (N-O) A very few multi-ciliated cells in the AIJ expressed FOXA2. (P-R) No FOXA2 expression in the ISM and UTJ. N=8 oviducts. (S-W) Luminal morphology and multi-ciliated cell distribution in the marmoset fallopian tube. (S-U) The fimbriae and AMP showed complex luminal epithelial folds. (U) Dispersed AT+ve multi-ciliated cells in the proximal AMP. (V) In the ISM, the lumen size is smaller, with thicker stromal and muscularis layer. Less complex folds (W) The fallopian tube-uterus junction. The lumen size is larger. Smooth luminal surface lined with epithelium. Gland-like structures were noted, with no multi-ciliated cells. (X-g) PAX8 and SOX17 expression pattern along the marmoset fallopian tube. (X-Z,c-e) PAX8 and SOX17 were expressed in non-ciliated secretory cells in the INF and AMP. In the proximal AMP, the proportion of AT+ve multi-ciliated cells were low but they were PAX8-ve (Z) and SOX17-ve (e). (a,f) In the ISM, all epithelial cells were PAX8+ve (a) and SOX17+ve (f). (b,g) In the tube-uterine junction, all epithelial cells were PAX8+ve (b) and SOX17+ve (g) including gland-like structures. N=3 marmosets. Scale bars = 20 µm.

We wondered whether these distal/proximal regionalities were evolutionarily conserved in other mammalian species, particularly in primates. The marmoset (*Callithrix jacchus*) is a non-human primate that has recently acquired a lot of attention because of its potential to be used as a genetic model for human diseases (Okano et al., 2012). The adult marmoset female reproductive tract is anatomically similar to that of the human— uncoiled, bowed fallopian tubes (FTs) extending from a single fused uterus (Cui & Matthews, 1994) into finger-like projections, called fimbriae, that encompass the ovary at the distal end (Suppl. Fig.5A,B, N=3 marmosets). As expected, the proportion of multi-ciliated cells was higher in the distal fimbriae and AMP relative to the proximal ISM (Fig.4S-V) while no multi-ciliated cells were observed in the fallopian tube (FT)-uterus junction (Fig.4W). In the epithelium of the fimbriae and AMP, PAX8 and SOX17 were expressed in non-ciliated secretory cells (Fig.4X-Z,c-e) while in the ISM, both PAX8 and SOX17 were uniformly expressed in both multi-ciliated and non-ciliated secretory cells (Fig.4a,f). All epithelial cells in the FT-Uterus junction expressed PAX8 and SOX17 (Fig.4b,g). These results indicate that distinct epithelial populations in the distal/proximal regions of the fallopian tube epithelium are evolutionarily conserved between mice and primates.

### A sharp boundary of distal and proximal cell populations between Turn 2 and 3

A sharp reduction in the proportion of multi-ciliated cells was observed between Turn 2 and 3 (Fig.3A,B). We found that the change in distribution pattern of PAX8+ve cells was complementary to the change in distribution pattern of FLTP-H2B-Venus+ve multi-ciliated cells (Fig.5A-H). The distal epithelium with high-proportion of FLTP-H2B-Venus+ cells showed low-proportion of PAX8+ve cells, while the proximal epithelium with low-proportion of FLTP-H2B-Venus+ve cells showed uniform PAX8+ve cells with a straight boundary (Fig.5B-D). In a transverse section, the folds with dispersed PAX8+ve cells showed high-proportion of FLTP-H2B-Venus+ve cells where their expression was mutually exclusive. On the other hand, the folds with uniform PAX8+ve cells showed dispersed FLTP-H2B-Venus+ve cells where they were co-expressed (Fig.5E-H). The distribution pattern of SOX17 was identical to that of PAX8+ve cells (Fig.5I,J). This boundary was always bevelled, with the proximal population extending distally on the mesosalpinx side, located on the inner side of Turn 2-3 (Fig.5K,L).

**Figure 5:**
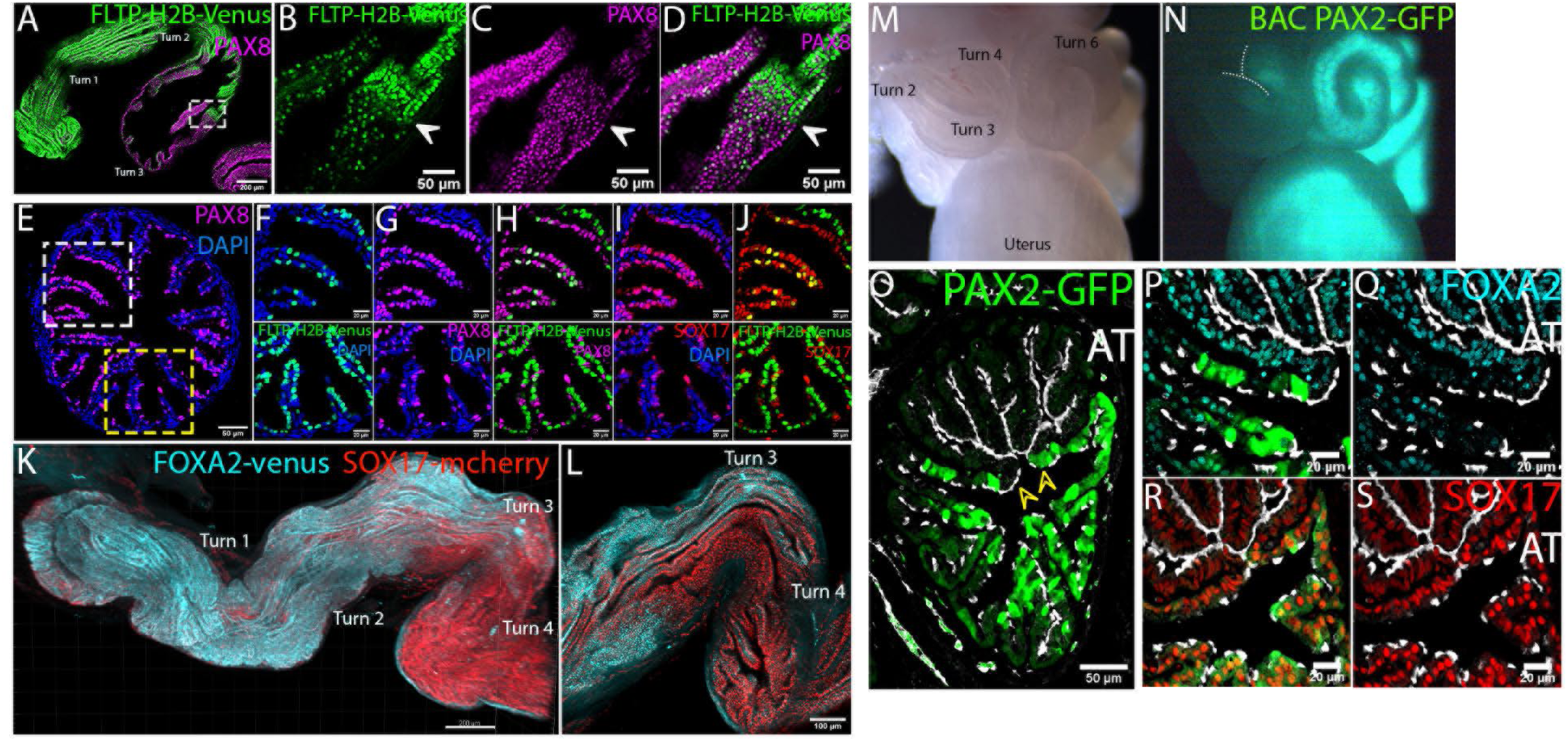
Sharp boundary of distal and proximal epithelial populations in the AIJ. (A-D) The sudden reduction of FLTP-H2B-Venus+ve cells coincides the uniform expression of PAX8. Scale bar = 200 µm. (B) A boundary of the high-proportion of FLTP-H2B-Venus+ve cells in the distal epithelium with the low-proportion of FLTP-H2B-Venus+ve cells in the proximal epithelium. (C) A boundary of the distal epithelium that PAX8 is specifically expressed in secretory cells with the proximal epithelium that PAX8 is uniformly expressed. (D) These two boundaries coincide (white arrowheads). Scale bars = 50 µm. (E) A transverse section of the boundary area. The folds with uniform PAX8+ve cells (white dotted box) and the folds with dispersed PAX8+ve cells (yellow dotted box). Scale bar = 50 µm. (F-H) (top panels) The folds with uniform PAX8+ve cells have dispersed FLTP-H2B-Venus+ve cells. PAX8 and FLTP-H2B-Venus are co-expressed. (bottom panels) The folds with dispersed PAX8+ve cells have high-proportion of FLTP-H2B-Venus+ve cells. PAX8 and FLTP-H2B-Venus are mutually exclusive. Scale bars = 20 µm. (I,J) SOX17 expression is identical to PAX8. (K) 3D distribution pattern of FOXA2-venus and SOX17-mcherry cells. Their boundary of distinct distal/proximal distribution patterns between Turn 2 and 3. Maximum projection. Scale bar = 200 µm. (L) Single optical section. The proximal epithelium with uniform SOX17+ve cells extended distally on the mesosalpinx side, the inner side of Turn 3, forming a bevelled boundary. N=4 mice. Scale bar = 100 µm. (M,N) Proximally restricted *Pax2*-GFP expression. GFP expression boundary around Turn 2-3. (O) Transverse section showing the separate distribution of *Pax2*-GFP+ve and GFP-ve cells. A couple of folds show GFP+ve cells on one side and GFP-ve cells in the other side of the fold (yellow arrowheads). The GFP+ve and GFP-ve epithelia opposed at the ridge of the fold. The GFP-ve epithelium had fewer multi-ciliated cells. Scale bar = 50 µm. (P,Q) Low proportion of FOXA2+ve cells in the *Pax2*-GFP+ve epithelium. (R,S) Uniform expression of SOX17in the *Pax2*-GFP+ve epithelium. N=4 mice. Scale bars = 20 µm.

In our parallel study, we demonstrated that PAX2 expression is restricted in the proximal oviductal epithelial cells (Ford et al., 2020). Using BAC *Pax2-GFP* mice (Pfeffer et al., 2002), the boundary of the proximal *Pax2*-GFP positive (*Pax2*-GFP+ve) and distal *Pax2*-GFP negative (*Pax2*-GFP-ve) cells was visualized between Turn 2 and 3, similar to the other boundaries above (Fig.5M,N). In a transverse section at the boundary, we observed separation of distal *Pax2*-GFP-ve cells and proximal *Pax2*-GFP+ve cells (Fig.5O). A couple of folds at this boundary showed *Pax2*-GFP+ve cells on one side and *Pax2*-GFP-ve cells on the other side of the same fold, opposed at the ridge of the fold. The boundary of the distinct distribution patterns of FOXA2+ve cells (Fig.5P,Q) and SOX17+ve cells (Fig.5R,S) also coincided. Therefore, all boundaries of the distinct distal/proximal expression patterns of PAX8, FOXA2, SOX17 and PAX2 coincided with the distinct distal/proximal distribution pattern of multi-ciliated cells between Turn 2-3. This supports the conclusion of our parallel study that the distal/proximal epithelial cells are separately maintained, distinct lineages (Ford et al., 2020).

### Proximally extended WT1 expression generates a distinct population in the distal PAX2 positive cells of the AIJ

WT1 is uniquely expressed in the distal luminal epithelial cells (Ford et al., 2020; Fig.6A-F). Interestingly, however, the boundary of WT1+ve and WT1-ve cells was shifted proximally to around Turn 6, beyond the sharp boundary mentioned above between Turn 2 and 3 (Fig.6G,J). WT1 was not restricted in the *Pax2*-GFP-ve distal cell population but also expressed in the *Pax2*-GFP+ve cell population of the AIJ (Fig.6B,C,G,H). The proportion of WT1+ve cells gradually decreased within the AIJ (Fig.6M, Suppl. Fig.6; 90% and 53% in distal and proximal AIJ, respectively). Interestingly, in the proximal AIJ between Turn 5 and 5.5, WT1+ve cells were predominantly non-ciliated/secretory cells, lining the short-broken longitudinal folds located on the anti-mesosalpinx side (Fig.3F and 6K,M). On the mesosalpinx side, WT1+ve cells were absent from transverse folds with ISM-like localization of FLTP-H2B-Venus+ve cells in trenches. This suggested that the proximal isthmic transverse folds extended distally on the mesosalpinx side into the area of WT1+ve short-broken longitudinal folds, forming a beveled boundary of AIJ-ISM. 3-4 patches of WT1+ve cells were observed in the distal ISM (Fig.6J,L). No WT1+ve cells were found in the proximal ISM (7% and 0% in distal to proximal ISM, respectively; Fig. 6D-F, M).

**Figure 6:**
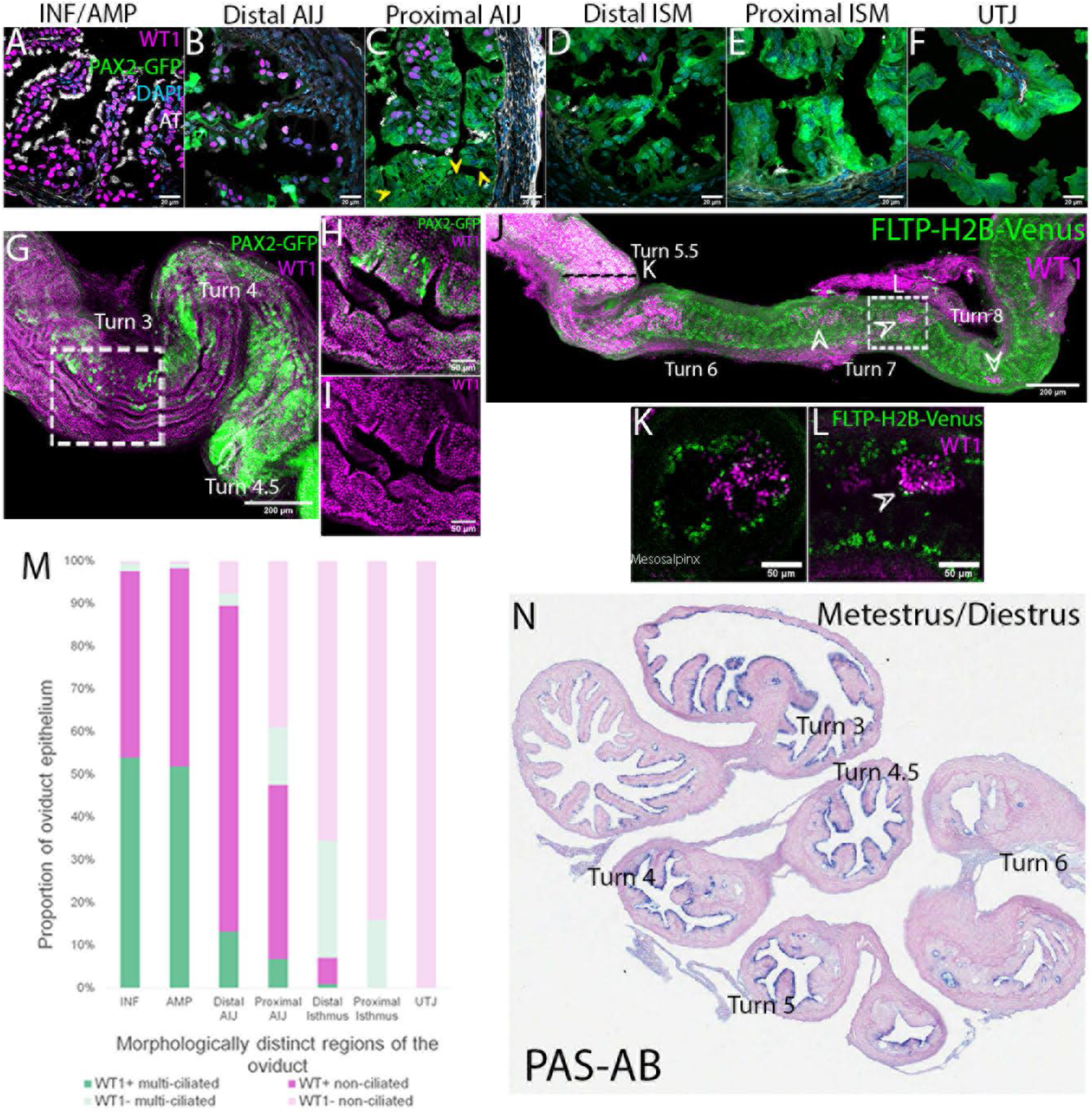
Boundary of WT1 expression extends into the proximal AIJ. (A-F) Distribution of WT1+ve and *Pax2*-GFP+ve cells in the oviduct epithelium. (A) INF/AMP. Uniform WT1+ve cells with no *Pax2*-GFP expression in the epithelium. (B,C) Distal and Proximal AIJ. Co-expression of WT1 and *Pax2*-GFP. Some *Pax2*-GFP+ve cells are WT1-ve in the proximal AIJ (yellow arrowheads). (D) Distal ISM. (E) Proximal ISM. (F) UTJ. Scale bar=20um. (G) WT1 expression extended into the proximal *Pax2*-GFP+ve population. Scale bar = 200 µm. (H,I) WT1 and PAX2 were co-expressed but the level of WT1 was lower in the *Pax2*-GFP+ve cells. N=4 oviducts. Scale bar = 50 µm. (J) WT1 expression boundary was located around Turn 6 with a few patches of WT1+ve cells in the distal ISM (white arrows). Scale bar = 200 µm. (K) Orthogonal view marked as the black dotted line in (J). WT1+ve cells were distributed on the folds on the anti-mesosalpinx side. The opposing mesosalpinx side had FLTP-H2B-Venus+ve cells in the trenches of transverse folds. They were not co-expressed. (L) Single optical section of the area indicated by a white dotted box in (J). Patches of coherent WT1+ve cells on the folds. WT1+ve cells were FLTP-H2B-Venus-ve. Scale bar = 50 µm. (M) Proportions of WT1+ve cells along the oviduct epithelium. Quantification was performed on transverse sections like A-F. (N) PAS-AB staining of the mouse oviduct during met/diestrus. Dark alcian blue staining was noted in only the AIJ epithelium.

Visualization of neutral-acid mucin secretion by PAS-AB staining showed dense alcian blue staining, indicating acid mucus secretion, only in the AIJ epithelium during the metestrus/diestrus stages, while no staining was noted in the AMP and ISM (Fig.6N; N=3 mice). In the estrus stage, no alcian blue staining was observed; some PAS staining, indicating neutral mucus secretion, was detected in the uterine glands (Suppl. Fig.7). Taken together, our results indicated that the AIJ is different from the AMP and ISM in luminal morphology, multi-ciliated cell distribution, WT1 expression and secretion regulation.

### Oocytes and preimplantation embryos travel along the oviduct during fertilization and preimplantation development

It is known that fertilization takes place in the AMP. However, it is still unknown how the oocyte and sperm entering from the opposite ends of the oviduct can meet consistently at the AMP. Following ovulation, the AMP was swollen and filled with a clear serous fluid (Fig.7A,B). The oocyte-cumulus complexes were located around Turn 2, within the AMP (Fig.7B,C) encompassing the boundary of the two distal and proximal epithelial populations, as visualized by the sudden decrease in multi-ciliated cell proportion (Fig.7D). Interestingly, the swollen AMP maintained its shape even after the oviduct was detached from the uterus and the ovary, suggesting that both ends of the AMP were closed with no fluid leakage (Fig.7A). Both ends of the swollen AMP appeared physically constricted to close the lumen (Fig.7E). When we injected trypan blue solution into the swollen AMP, the solution easily diffused into the AMP lumen but entered neither the AIJ nor the ovarian bursa (Fig.7F,G; N=3 mice). In contrast to this, when we tried to inject into the ISM, a very small amount of the dye solution could be injected, the solution did not distribute in either direction within the ISM luminal space (Fig.7F,H). When we flushed the oviduct from the INF, a very thick white mucus was flushed out from the oviduct (Fig.7I,J). This suggests that, in addition to the physical constriction, the proximal region of the oviduct was plugged during ovulation with thick mucus blocking any fluid flow and oocyte movement into the ISM.

Although the oviduct is known to be the place where preimplantation development occurs (Kölle et al., 2020; Moore et al., 2019), it is unknown whether the embryos are freely floating in the oviduct lumen. At 0.75 dpc, zygotes were located within the AIJ (Fig.7K,L), suggesting that the thick mucus was removed from the region. At 1.5 dpc, a 2-cell stage embryo was in the proximal ISM close to the UTJ (Fig.7M,N), appearing free floating in the lumen with occasional ISM contractions. Interestingly, at 2.5 dpc, a single queue of morula embryos halted in the UTJ (Fig.7O-Q), within the compartments found at the oviduct-uterus junction (Fig.2P).

**Figure 7:**
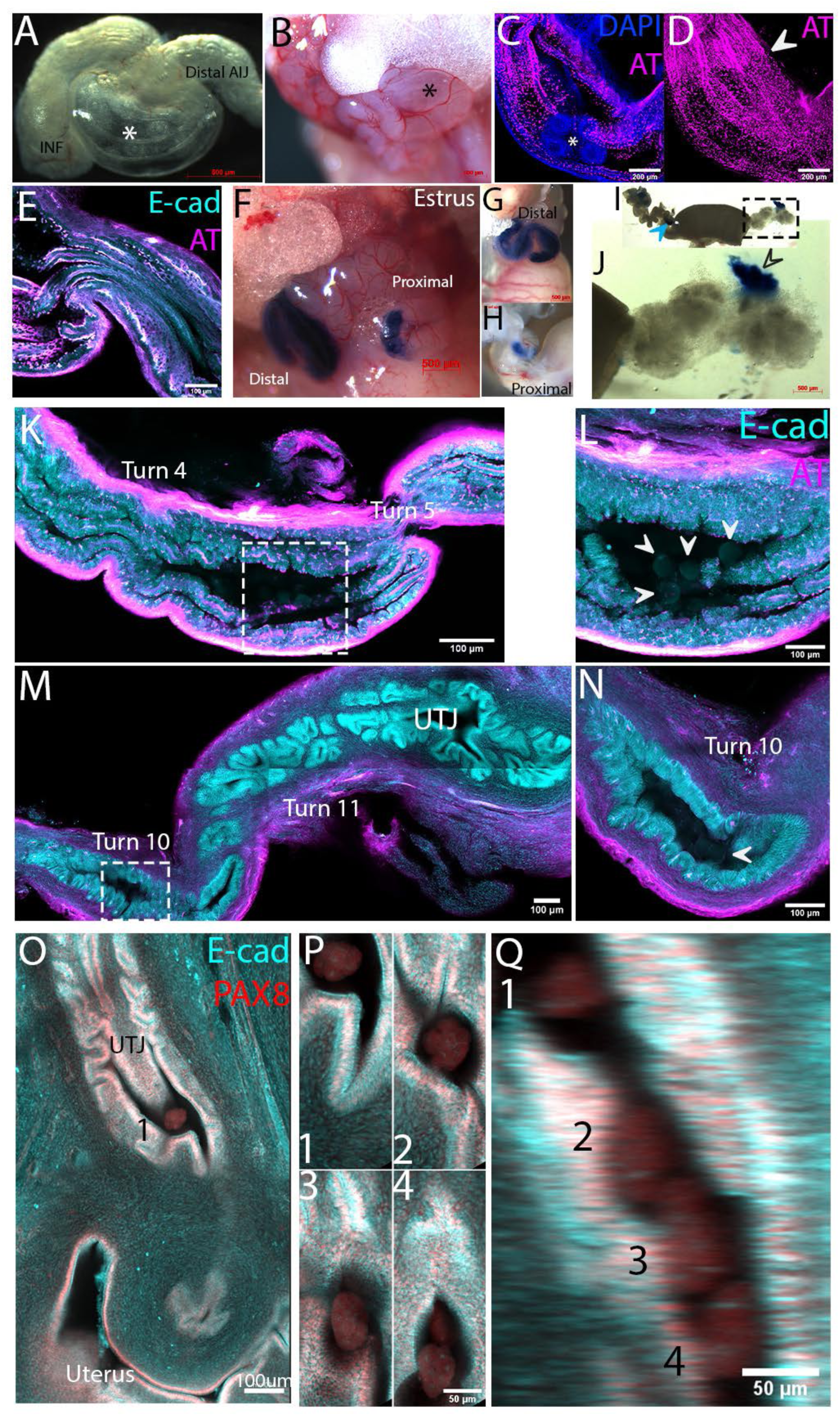
Preimplantation embryo travel in oviduct luminal space. (A) Prominent swollen AMP after ovulation (black asterisk). (B) A swollen AMP after dissection. Several oocyte-cumulous complexes are visible (white asterisk). Scale bar=500um. (C) A swollen AMP containing 0.5 dpc embryos (white asterisk). Single optical section. (D) Maximum projection of AT staining of (C). The boundary of the distal/proximal populations (white arrow). Scale bar = 200 µm. (E) Single optical section showing the AIJ side of a swollen AMP. The lumen was closed by lumen constriction. Scale bar = 100 µm. (F-H) Injection of trypan blue solution. (G) In the AMP and INF, trypan blue solution spread evenly but did not into the AIJ or the ovarian bursa. (H) In the ISM, only a small amount of trypan blue solution could be injected. The blue dye did not spread but stayed at the injection site. (I) A thick viscous mucus flushed out from the oviduct. A blue arrow indicates the dye injection site in the ISM. (J) Higher magnification of the mucus. A part of the mucus has blue dye (black arrowhead) indicating this position of the mucus was at the dye injection site in the ISM. Scale bar = 500 µm. (K) 0.75 dpc 1-cell stage embryos in the AIJ around Turn5. Characteristic broken-longitudinal folds are observed, with sparsely distributed multi-ciliated cells. Four zygotes are in the area marked with a dotted box. (L) Higher magnification of the area marked in (K). Floating zygotes (white arrowheads). This suggests that the mucus was cleared in this area by this time point. (M) 1.5dpc. a 2-cell stage embryo in the proximal ISM around Turn10. A 2-cell embryo is in the area marked in a dotted box. (N) Higher magnification of the area marked in (M). A 2-cell embryo (white arrowhead). Scale bar = 100 µm. (O-Q) 2.5dpc. Four morula embryos forming a queue in the compartments of the UTJ. (O) Compartments were located at the junction of the oviduct and uterus. The first embryo and uterine lumen are shown. Scale bar=100um. (P) Four embryos located at different z-positions along the UTJ. (Q) YZ orthogonal view. Four morula embryos formed a queue. N=3 mice. Scale bar = 50 µm.

## DISCUSSION

The oviduct is an essential tissue for reproduction in mammals, however, its anatomical complexity and cellular heterogeneity have not been fully appreciated. Using modern imaging techniques and mouse fluorescence reporter lines, we identified previously unrecognized anatomical structures and cellular heterogeneity in the oviduct luminal epithelium (Fig.8A). We showed that basic turning points were associated with distinct luminal morphologies and epithelial populations. Previous studies often incorrectly compare different regions of the oviduct in traditional 2D sections, due to lack of landmarks for the coiling oviduct. Therefore, the turning points can be used as reliable landmarks to help precise description and comparison possible within the oviduct.

How are the turning points associated with luminal regionalities? We speculate that the oviduct employs a mechanism similar to that of the intestine. The intestine, a well-known looped organ, has a consistent looping pattern with occasional variation due to contractile movement. Its looping points are tightly linked with intestine regionalities and artery access points (Soffers et al., 2015). Prior to looping, the artery network reaches specific sites of the intestine via the mesentery, a membranous organ connecting the entire intestinal tube. The looping morphogenesis follows the pre-existing regionalities and artery network. Mechanical contributions of the mesentery are crucial for this morphogenesis (Huycke et al., 2019; Nerurkar et al., 2017). This is a developmentally controlled process with minimum variation. The oviduct-mesosalpinx structural relationship and the sequence of the developmental events appears similar to that of the intestine-mesentery.

The three luminal transitions at the AMP-AIJ, AIJ-ISM and ISM-UTJ were always beveled, with proximal characteristics extending distally on the mesosalpinx side, i.e. the inner side of the turns. We also noticed that the distal-most AMP-AIJ transition was wider than the ISM-UTJ. This pattern looks similar to the starting position difference between inner and outer tracks of 200- or 400-meter sprints, as the goal position is aligned at the opposite end. We speculate that the luminal regionalities in the oviduct are specified first with blunt boundaries (Fig.8B). Subsequently, these regionalities guide morphogenetic coiling events. Since oviduct coiling is the consequence of multiple turns toward the mesosalpinx side, the distance gap between the inner/outer epithelia makes the boundaries beveled. Longer the distance from the aligned point, the gap becomes wider. This model is testable in the mouse line with constitutive Notch activation in oviductal stromal cells, which has a deficiency in oviduct coiling and normal luminal morphology in the adult (Ferguson et al., 2016).

**Figure 8:**
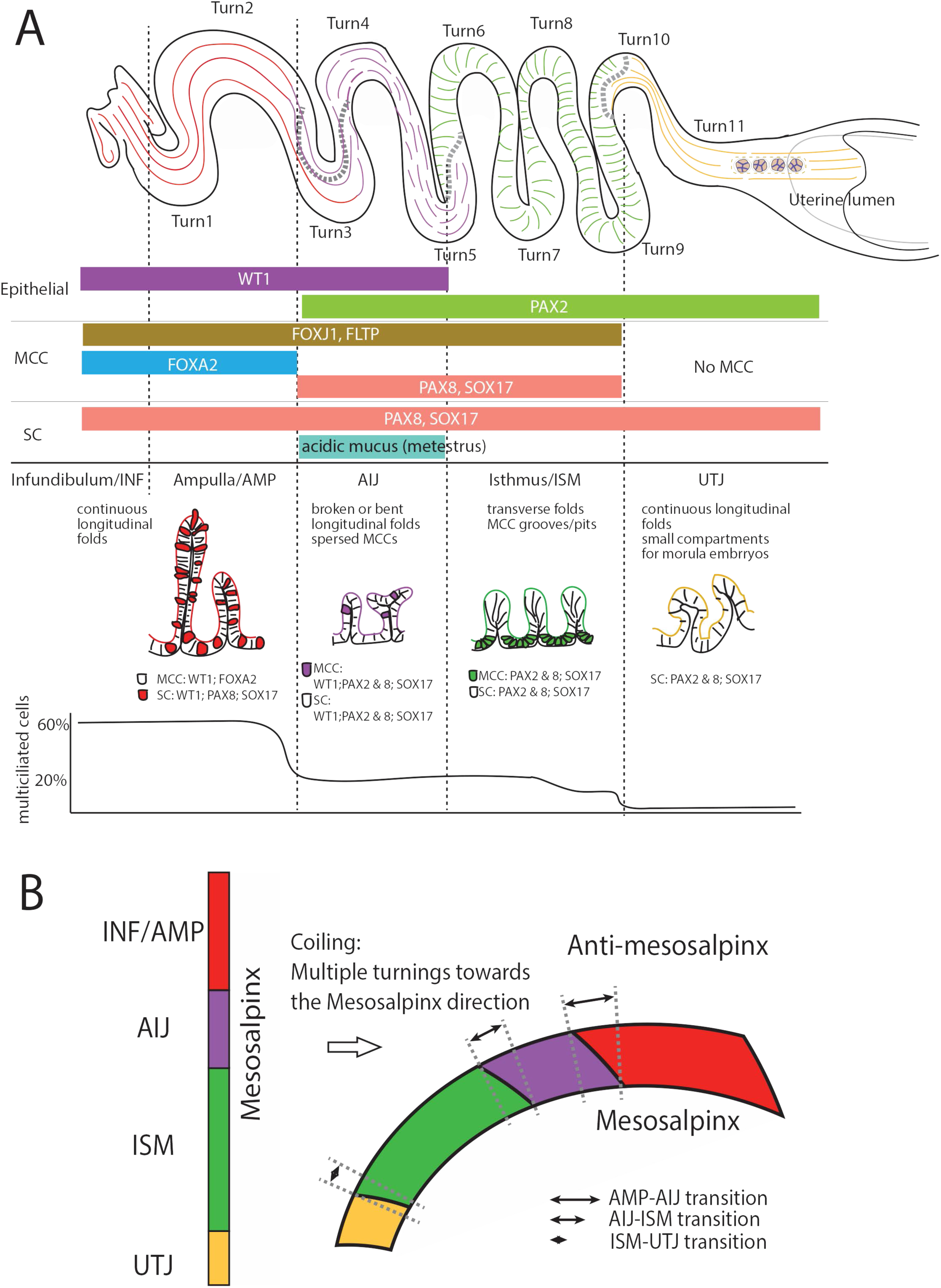
Models of anatomical and cellular heterogeneity of the oviductal luminal epithelium and formation of boundary beveling. (A) Anatomical and cellular heterogeneity of the oviductal luminal epithelium. Turning points are linked with luminal morphological transitions. Five distinct regionalities; INF, AMP, AIJ, ISM and UTJ, can be defined by luminal epithelial fold morphology, multi-ciliated cell distribution pattern, transcriptional factor expression pattern and mucus secretion. The distribution of multi-ciliated cells did not form a gradient but had three distinct proportions: high (INF/AMP), low (AIJ/ISM) and none (UTJ). Small compartments for morula embryos in the UTJ are also shown. A model for formation of boundary beveling. (left) The blunt boundaries are specified before coiling. (right) Coiling, multiple turns toward the mesosalpinx side, causes the distance gap between the inner and outer side of the tube. The base of the oviduct is less affected by coiling but further away from the base, the gap becomes wider.

In the distal region, the primary role of multi-ciliated cells is to support movement of the cumulus oocyte complex (COC) (Talbot et al., 1999). On the other hand, multi-ciliated cells in the ISM formed pits and grooves. It is not clear if they can create a fluid flow in the luminal space to support preimplantation embryo movement since they were located in the trenches of transverse luminal folds with multi-cilia projecting into focal points. Interestingly, the stripe pattern of multi-cilia in the ISM is reminiscent of the immobilized sperm localization pattern (La Spina et al., 2016; Muro et al., 2016). SEM studies of bull sperm indicate attachment of uncapacitated sperm to multi-ciliated cells of the oviduct (Lefebvre et al., 1995). Studies in various species including mice allude to the presence of sperm attachment sites in the ISM, usually called sperm reservoirs (Maillo et al., 2016; Sasanami et al., 2013; S. S. Suarez, 2001, 2002, 2016). The multi-ciliated cell clusters we identified in the ISM could function as sperm reservoirs, playing important roles in sperm viability, hyperactivation and capacitation (Maillo et al., 2016; Suarez, 2016).

Classic transmission electron microscopic (TEM) studies in various mammalian species suggest that the distal and proximal oviduct epithelial cells have distinct secretary functions (Abe, 1996; Abe & Oikawa, 1991; Lauschová, 1999, 2003; Murray, 1997). High to moderate electron dense secretory granules in the INF and AMP suggest proteinaceous, serous secretion. Predominantly low electron dense secretory granules suggest mucus containing secretory granules in the ISM (Abe, 1996; Abe & Oikawa, 1991; Murray, 1997). Our results were consistent with these classic observations. In the distal region, a clear serous fluid was observed in the swollen AMP during ovulation. In the ISM, a highly viscous mucus was observed transiently during estrus stage. Similar isthmic mucus accumulation during the estrous stage is noted in macaques, rabbits, and humans (Hunter et al., 2011; Jansen, 1978, 1995). It has been shown that very few sperm reach the point of fertilization in the AMP even 4 hours after mating (La Spina et al., 2016; Muro et al., 2016). Additionally, the sperm collected from several mutant mouse lines, including ADAM3 KO mice, present with no clear motility defect but fail to enter the oviduct via the UTJ (Fujihara et al., 2019; Yamaguchi et al., 2009). It is interesting to speculate that those genes are necessary for sperm to penetrate the highly viscoelastic mucus plug in the ISM. This mucus plug likely prevents polyspermy and entry of unhealthy sperm into the AMP (Hunter, 2012; Hunter et al., 2011). Various secretion molecules like anti-bacterial and fungal peptides are also secreted from the oviductal luminal epithelial cells in a hormonal cycle dependent manner (Barton et al., 2020; Ghersevich et al., 2015; Li & Winuthayanon, 2017; Winuthayanon et al., 2015). How these secretions are regulated will be very important to understand gynecological diseases and infertility.

A sharp boundary of the distal and proximal populations was located between Turn 2 and 3. Interestingly, the boundary of WT1 expression did not coincide with this boundary but was shifted proximally. This created a unique cell population in the AIJ that expressed WT1 in the proximal PAX2+ve population (Fig.8A). The unique luminal AIJ morphology is briefly reported previously (Agduhr, 1927; Barton et al., 2020; Burton et al., 2015). In addition to WT1 expression, we found three other unique characteristics of the AIJ. First, longitudinal folds in the AIJ were not continuous but had multiple breaking points and convolutions. Second, PAX8+ve multi-ciliated cells on the folds were sparsely distributed. Third, specific alcian blue staining during metestrus/diestrus stages. Taken together, we propose that the AIJ is another regionality, with distinct functions and unique gene expression.

Mouse preimplantation embryos stay in the oviduct for 3 days before implanting into the uterine lining (Kölle et al., 2020; Moore et al., 2019). By E2.5, around the 8-cell stage, they moved into a previously unrecognized compartment in the UTJ, where the oviduct connects into the uterus. In contrast to earlier stages where the embryos appeared free-floating in the ISM, morula embryos formed a queue in the UTJ, as if waiting for the entry to the uterus. It will be very interesting to determine what controls the timing of embryo entry into the uterus.

The intimate relationship between the oviduct epithelial cells and gametes/preimplantation embryos is essential for successful pregnancy. Our work provides a foundation to understand the oviduct luminal environment and homeostasis in reproduction.

## METHODS AND MATERIALS

### Animals

All animal work was performed in accordance with institutional guidelines and was approved by the Faculty of Medicine Animal Care Committee (AUP #7843), it was undertaken at the Goodman Cancer Research Centre animal facility. *Pax2-GFP Bac* (Pfeffer et al., 2002) mice were kindly provided by Dr. Maxime Bouchard (McGill). *Fltp*-*H2B-Venus* (Gegg et al., 2014), *FoxA2-venus* (Burtscher et al., 2013), *Sox17-mCherry* mice (Burtscher et al., 2012) mice were generated by Dr. Heiko Lickert (IDR Munich, Germany). Adult mice used in this study were 2-4 months of age, estrous stages were analyzed using vaginal smears stained with crystal violet (McLean et al., 2012). Necropsy specimens of adult female marmoset reproductive tracts were kindly provided by Dr. Jim Gourdon (CMARC, McGill) and Dr. Keith Murai (McGill).

### Whole mount immunostaining, tissue clearing and 3D confocal imaging

After euthanasia, mouse female reproductive tracts were collected and straightened by removing the mesosalpinx. The straightened oviduct was fixed with DMSO: methanol (1: 4) and cut into 3-4 pieces prior to placing at −20°C overnight. The antibody staining protocol was as described in Arora et al., 2016. BABB cleared samples were placed on a #1.5 coverslip (Fisher Scientific, 12-545F) with 10-15ul BABB prior to imaging using the 10X objective (NA 0.30) on LSM 800 or 710 (Zeiss). Section interval for 3D confocal imaging was 4.32 µm. Usually 40-90 optical sections were taken.

### Antibodies

Primary antibodies (1/250 dilution): anti-PAX8 (Proteintech, 10336-1-AP), anti-GFP (Abcam, ab13970), anti-mCherry (Abcam, ab213511), anti-WT1 (Abcam, ab89901), anti-acetylated tubulin (Sigma, T7451), anti-Sox17 (R&D, AF1924), anti-FoxA2 (Cell Signaling, 3143), and anti-E-cadherin (Invitrogen, 13-1900). Secondary antibodies (1/450 dilution): Alexa Fluor (AF) anti-rabbit 555 (Invitrogen, A31572), AF anti-rabbit 649 (Invitrogen, A21245), AF anti-rabbit 488 (Invitrogen, A21206), AF anti-mouse 649 (Invitrogen, A32787), AF anti-mouse 488 (Invitrogen, A21202), AF anti-goat 568 (Invitrogen, A11057), AF anti-rat 488 (Invitrogen, A21208), anti-chicken 488 (Sigma, SAB4600031), DAPI/Hoechst 33342 (Thermo Fisher, 62249), Alexa Fluor 488 phalloidin (Lifetech, A12379), and Alexa Fluor 635 phalloidin (Lifetech, A34054).

### Trypan blue injection and oviduct flushing

Estrous stage pseudo-pregnant females were anesthetized. The female reproductive tract was exposed either dorsally or ventrally. Under a dissection microscope, trypan blue solution (STEMCELL Technologies, #07050) was injected into the swollen AMP or the ISM using a glass needle. Mice were euthanized, the reproductive tracts were dissected out and straightened slightly.

### Image analysis, cell counts, and statistics

3D confocal images were stitched using ImarisStitcher and visualized using FIJI and Imaris. 3D rendering was performed using surfaces function and distance transformation (Outside a Surface Object) MATLAB XTension, followed by manual background removal on Imaris. 2D sections were visualized using FIJI and Zen. For optical projection, Maximum/Average/Sum Slices projection in FIJI was used. Measurements and epithelial cell counting were performed manually. Mann-Whitney two-tailed test was performed to gauge significance. In some images of Fig.2, PAX8 immunostaining and FLTP-H2B-Venus were overlaid with a single color (cyan) to visualize the epithelial layer in the oviduct, indicated as Epithelium.

## Acknowledgement

We thank the McGill Goodman Cancer Research Centre Histology, the McGill Advanced Bioimaging Facility (ABIF) and the McGill integrated core of animal modeling (MICAM) for technical support. We thank Dr. Maxime Bouchard for the Pax2-GFP mouse line. We also thank Drs Keith Murai and Jim Gourdon for marmoset necropsy specimens. This work was supported by Canadian Cancer Society (CCS) Innovation grant (Haladner Memorial Foundation #704793), CCS i2I grant (# 706320) and Cancer Research Society Operation Grant (#23237). K.H. was supported by CRRD, Alexander McFee, and Rolande & Marcel Gosselin studentships. M.F. was supported by Canderel, CRRD and FRQS postdoc fellowships. K.T. was supported by MICRTP and Canderel studentships.

## FIGURE LEGENDS

**Video 1: A beveled flat surface of the ostium with radially asymmetric infundibular exterior folds**

The video showing a maximum projection of E-cadherin staining of the INF, followed by optical sections showing exterior and luminal folds (magenta). The luminal shape (gray) and exterior folds (blue) were rendered using Imaris. The relatively flat surface of the beveled end the oviduct with the ostium at the center and a side slit. The exterior folds on the slit formed an overhang. The back side of the ostium had flattened exterior folds.

**Video 2: Unique luminal compartments at the base of the oviduct**.

The video showing the connection of UTJ to the uterine lumen. The oviductal lumen approached from the mesosalpinx side connecting to the uterine lumen through the colliculus tubarius. At the junction of the oviduct and uterus, small luminal compartments created by simultaneous breakpoints in several longitudinal folds.

**Supplementary Figure 1: Wire models of oviduct coiling patterns**

(A) A wire model of a right oviduct. (B) The TDE cleared oviduct modeled in (A). (C) A wire model of another right oviduct. (D) The TDE cleared oviduct modeled in (C). (E) A wire model of a third right oviduct. (F) The TDE cleared oviduct modeled in (E).

**Supplementary Figure 2: Relative position of the ovary, INF and uterus in a 2D tissue section**.

H&E staining of a mouse female reproductive tract with the ovary. The INF is located inside of the ovarian bursa adjacent to the ovary. The exterior folds on the ovarian side are taller and complex than the opposite side. The mucosal epithelium of exterior folds was continuous with the epithelium lining the ovarian bursa and ovarian hilum (black arrows).

**Supplementary Figure 3: Transverse sections of distinct luminal fold patterns and multi-ciliated cell distribution**.

(A-G) Transverse sections of different areas of the oviduct. Top panels are F-actin staining. Bottom panels are FLTP-H2B-Venus and acetylated tubulin (AT) staining, both marking multi-ciliated cells. (A) INF, white arrowhead shows flattened exterior folds continuous with connective ligament. (B) AMP. (C) Distal AIJ. (D) Proximal AIJ. (E) Distal ISM. (F) Proximal ISM. (G) UTJ. The quantification of the proportions of FLTPH2B-GFP+ve cells was performed and presented in Figure 3G. N=4 mice.

**Supplementary Figure 4: Changes in proportions of PAX8+ve, SOX17+ve and FOXA2+ve cells in multi-ciliated and non-ciliated secretory cells along the mouse oviduct**.

(A) Proportion of PAX8+ve cells in multi-ciliated and secretory cells of total epithelial cells. (B) Proportion of PAX8+ve cells in respective cell types. (C) Proportion of SOX17+ve cells in multi-ciliated and secretory cells of total epithelial cells. (D) Proportion of SOX17+ve cells in respective cell types. (E) Proportion of FOXA2+ve cells in multi-ciliated and secretory cells of total epithelial cells. (F) Proportion of FOXA2+ve cells in respective cell types. N=8 oviducts. Quantification was performed on transverse tissue sections like Fig.4. Error bars indicate standard error. ****<0.0001, **<0.01, *<0.05.

**Supplementary Figure 5: Marmoset’s fallopian tube**

Marmoset fallopian tube. Prominent fimbriae, finger-like projections, are visible at the rostal end (arrow). No coiling. One bend (black asterisk) (B) H&E staining of fimbriae located adjacent to the ovary and ISM after the bend.

**Supplementary Figure 6: Proportion of WT1+ve cells in multi-ciliated and secretory cells**.

Proportion of WT1+ve cells in respective cell types. Quantification was performed on transverse tissue sections like Figure 6A-F. N=8 oviducts. Error bars indicate standard error. ****<0.0001, **<0.01, *<0.05.

**Supplementary Figure 7: PAS-AB staining in estrus stage**.

PAS-AB staining in the estrus stage. No alcian blue staining in the estrus stage mouse oviduct. Red PAS staining in uterine glands (black arrows).

