## Supplementary material for "Anatomical and cellular heterogeneity in the mouse oviduct-- its potential roles in reproduction and preimplantation development": Suppl figures

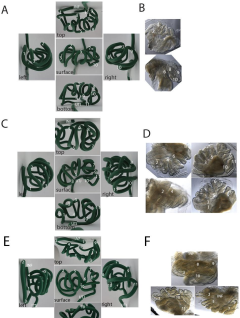


Harwalkar et al., 2020
Suppl. Fig. 1

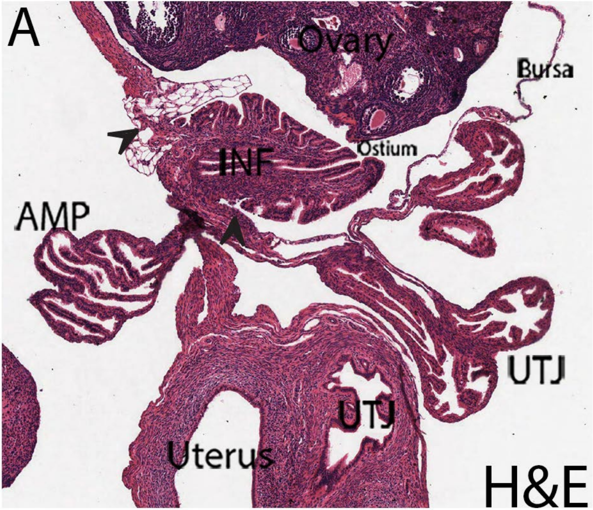


Suppl. Fig. 2
Harwalkar et al., 2020

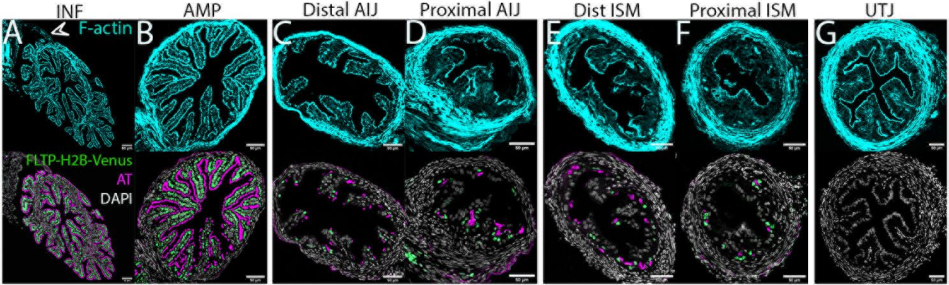


Suppl. Fig. 3
Harwalkar et al., 2020

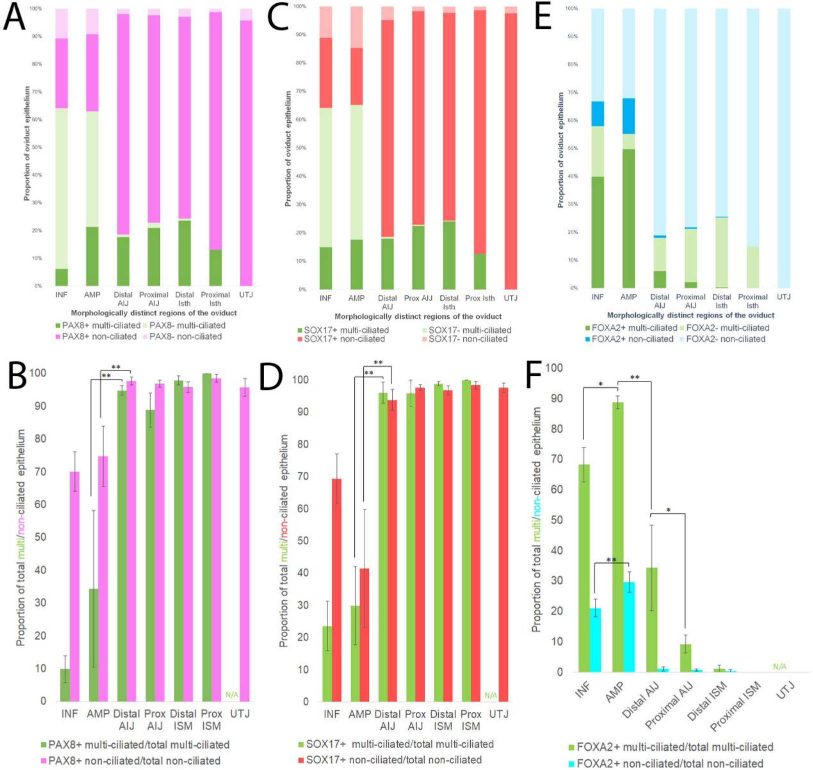


Suppl. Fig. 4
Harwalkar et al., 2020

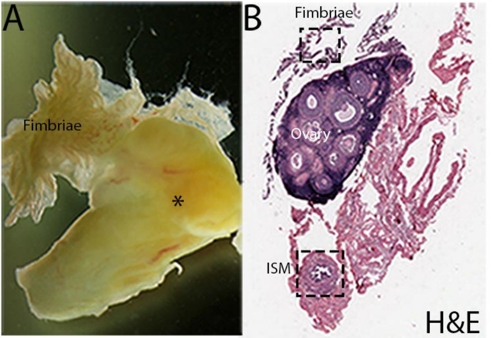


Suppl. Fig. 5
Harwalkar et al., 2020

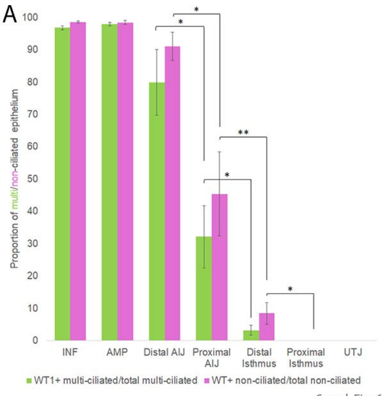


Suppl. Fig. 6
Harwalkar et al., 2020

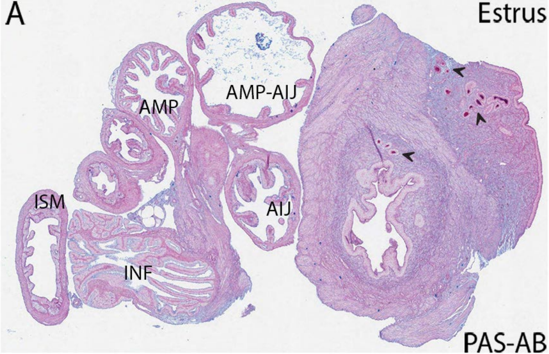


Suppl. Fig. 7
Harwalkar et al., 2020
